# Active mRNA degradation by EXD2 nuclease elicits recovery of transcription after genotoxic stress

**DOI:** 10.1101/2022.07.20.499545

**Authors:** Jérémy Sandoz, Max Cigrang, Philippe Catez, Lise-Marie Donnio, Clèmence Elly, Pietro Berico, Cathy Braun, Sergey Alekseev, Jean-Marc Egly, Guiseppina Mari-Giglia, Emmanuel Compe, Frédéric Coin

## Abstract

The transcriptional response to genotoxic stress involves gene expression arrest, followed by recovery of mRNA synthesis (RRS) after DNA repair. Using a small-scale RNA interference screen, we found that the lack of the EXD2 nuclease impaired RRS and decreased cell survival after UV irradiation, without affecting DNA repair. Overexpression of wild-type, but not nuclease-dead EXD2, restored RRS and cell survival. We observed that UV irradiation triggered recruitment of EXD2 to chromatin where the nuclease transiently interacts with RNA Polymerase II (RNAPII) to promote the degradation of nascent mRNAs synthesized at the time of genotoxic attack. Reconstitution of the EXD2-RNAPII partnership on a transcribed DNA template *in vitro* showed that EXD2 primarily interacts with an elongation-blocked RNAPII and efficiently digest mRNA. Overall, our data highlight a crucial new step in the transcriptional response to genotoxic attack in which EXD2 interacts with elongation-stalled RNAPII on chromatin to degrade the associated nascent mRNA, allowing transcription restart after DNA repair.

## Introduction

Cells are regularly exposed to endogenous and exogenous genotoxic attacks that induce damage in the DNA molecule (Hoeijmakers, 2009), (Vermeij et al., 2014). The generation of DNA damage can potentially challenge several fundamental cellular processes such as transcription or replication and can ultimately cause diseases such as cancer if not repaired (Kotsantis et al., 2018), (Lee and Young, 2013), (Jackson and Bartek, 2009). The identification of several protective mechanisms against genotoxic stress highlights the importance of maintaining genome integrity to ensure low mutation frequencies in the genome (Edifizi and Schumacher, 2015). One such mechanism, the nucleotide excision repair (NER) pathway, removes DNA adducts such as pyrimidine (6-4) pyrimidone (6-4PP) or cyclobutane pyrimidine dimers (CPD) that are produced by UV light (Sancar and Reardon, 2004), (Marteijn et al., 2014), (Scharer, 2013). Two NER sub-pathways co-exist in cells: global genome NER (GG-NER), which removes DNA damage from the entire genome, and transcription-coupled NER (TC-NER), which corrects lesions located on actively transcribed genes (van den Heuvel et al., 2021), (Ljungman and Lane, 2004), (Hanawalt and Spivak, 2008), (Marteijn et al., 2014). In GG-NER, the concerted action of XPC and/or DDB2-containing complexes enables detection of DNA damage in the genome, whereas in TC-NER, an actively transcribing RNA Polymerase II (RNAPII), which is stalled by a lesion, triggers efficient removal of cytotoxic damage (Fousteri and Mullenders, 2008), (Lans et al., 2019).

To protect the integrity of gene expression under genotoxic attack, cells undergo a transcription stress response that includes global inhibition of transcription occurring in two steps: rapid and local inhibition of elongation due to the stalling of RNAPII in front of transcription-blocking DNA damage (Lavigne et al., 2017) which is followed by a global inhibition of transcription initiation occurring on both damaged and undamaged genes (Gyenis et al., 2014), (Rockx et al., 2000). Recent evidence has shown that global inhibition takes place after degradation of the pool of RNAPII (Tufegdžić Vidaković et al., 2020), (Nakazawa et al., 2020). After DNA repair, cells recover transcription in an active process involving transcription and chromatin remodelling factors (Oksenych et al., 2013), (Mourgues et al., 2013), (Adam et al., 2013), (Dinant et al., 2013), (Geijer et al., 2021). Recovery of RNA synthesis (RRS) encompasses both the re-initiation of expression at the promoters of actively transcribed genes and the restart of RNAPII molecules already in elongation. Despite recent advances in our understanding of the transcriptional stress response to genotoxic attack, the actors and mechanisms responsible for RRS after DNA repair remain largely elusive. Finding new actors involved in RRS is therefore crucial to better understand this process at the molecular level and its role in genome stability.

Using an *in cellulo* reporter system coupled to a small-scale genetic screen, we unveil here that EXD2, a RNA/DNA nuclease previously shown to be involved in homologous recombination and in the replication fork protection pathway (Nieminuszczy et al., 2019)(Broderick et al., 2016), is essential for RRS after genotoxic attack. Cells lacking EXD2 or expressing a nuclease-dead version of the enzyme are unable to restore global RNAPII-dependent transcription after UV irradiation, resulting in decreased resistance to genotoxic attack. Mechanistically, we demonstrated that EXD2 is not involved in the removal of UV-induced photoproducts. Instead, UV irradiation provokes translocation of EXD2 to chromatin where it transiently interacts with RNAPII and promotes the degradation, during the recovery phase of transcription, of nascent mRNA being synthesized at the time of genotoxic attack. Using a reconstituted transcription system *in vitro*, we reconstructed the dynamic association of EXD2 to RNAPII on a transcribed DNA template and demonstrated that EXD2 preferentially interacts with an elongation-blocked RNAPII. In such system, the ribonuclease activity of purified EXD2 efficiently processes mRNA. These findings unveil a crucial role for EXD2 in the transcription stress response and are the first to assign a nuclear function to the ribonuclease activity of EXD2 by showing its involvement in the degradation of mRNA under synthesis at the time of genotoxic attack. This degradation is necessary for an efficient recovery of gene expression after DNA repair.

## Results

### UV-induced inhibition and recovery of transcription at a defined genomic locus

In order to identify novel factors required for transcription recovery after a genotoxic attack, we first sought to develop a sensitive assay to easily monitor transcription inhibition and recovery after UV irradiation. As shown in Supplemental Figure 1a, we used a doxycyclin (dox)-inducible transcription/translation reporter system integrated at a single site on genomic DNA in the human osteosarcoma U-2 OS cell line. This system allows visualization of the genomic locus, its nascent mRNA transcript (*CFP-SKL*) and protein product (CPF-SKL) (Janicki et al., 2004), (Shanbhag et al., 2010). After a 2-hour dox treatment we detected transcription of *CFP-SKL* in 80% of the cells (Supplemental Figure 1b). A 2-hour dox treatment followed by a recovery period in the absence of dox (1-to 4-hours) triggered an accumulation of CFP-SKL protein expression (Supplemental Figure 1c). The plasticity of this system also allowed us to measure the transcriptional activity of the cells in a specific time window after UV-irradiation. For this purpose, cells were irradiated with UV (30J/m^2^) and pulsed with dox for 2 hours at different times after genotoxic attack. Under these conditions, we noticed a strong inhibition of *CPF-SKL* mRNA expression at early times after UV irradiation (Supplemental Figure 1d, right panel, lanes 1 and 2). Interestingly, *CPF-SKL* mRNA expression recovered over time after irradiation (lane 3) and knockdown of the TC- and GG-NER factor XPA prevented this recovery (lanes 4-6 and left panel). CPF-SKL protein expression followed that of its mRNA with a strong inhibition early after irradiation and a progressive recovery that was completed 18 hours after irradiation (Supplemental Figure 1e). These data indicate that *CPF-SKL* expression recapitulates the rapid inhibition and progressive recovery of general transcription that is generally completed 20 hours after a genotoxic attack when DNA repair is efficient (Mayne and Lehmann, 1982), (Williamson et al., 2017).

### EXD2 is a critical factor for RRS

To explore the mechanism of transcription recovery after genotoxic stress, we used the reporter assay described above to conduct a small-scale genetic screen based on the transfection of siRNA pool targeting a dozen of enzymes, and/or their regulatory subunits, involved in transcription and/or DNA repair. Representative results of this screening are shown in Figure 1a. As expected, knockdown of the TC-NER factors CSA and CSB inhibited recovery of *CFP-SKL* expression. Interestingly, the best hit as potential effector of transcription recovery after UV irradiation was the 3’ to 5’ DNA/RNA exonuclease EXD2 (Figures 1a-b). Lack of recovery of *CFP-SKL* expression in the absence of EXD2 was confirmed by RT-qPCR (Figure 1c) and resulted in a dramatic inhibition (70%) of *de novo* CFP-SKL translation in time after UV-irradiation (Figure 1d, compare lanes 10-14 and 3-7). Of note, we observed that the accumulation of nascent *CFP-SKL* mRNA or *de novo* CFP-SKL protein was not impaired by the knockdown of EXD2 in the mock-treated cells (Figure 1b, compare panels a-b-c with j-k-l and Figure 1d, compare lane 2 and 9). These data suggest that EXD2 is required for RRS following a genotoxic stress.

**Figure 1:**
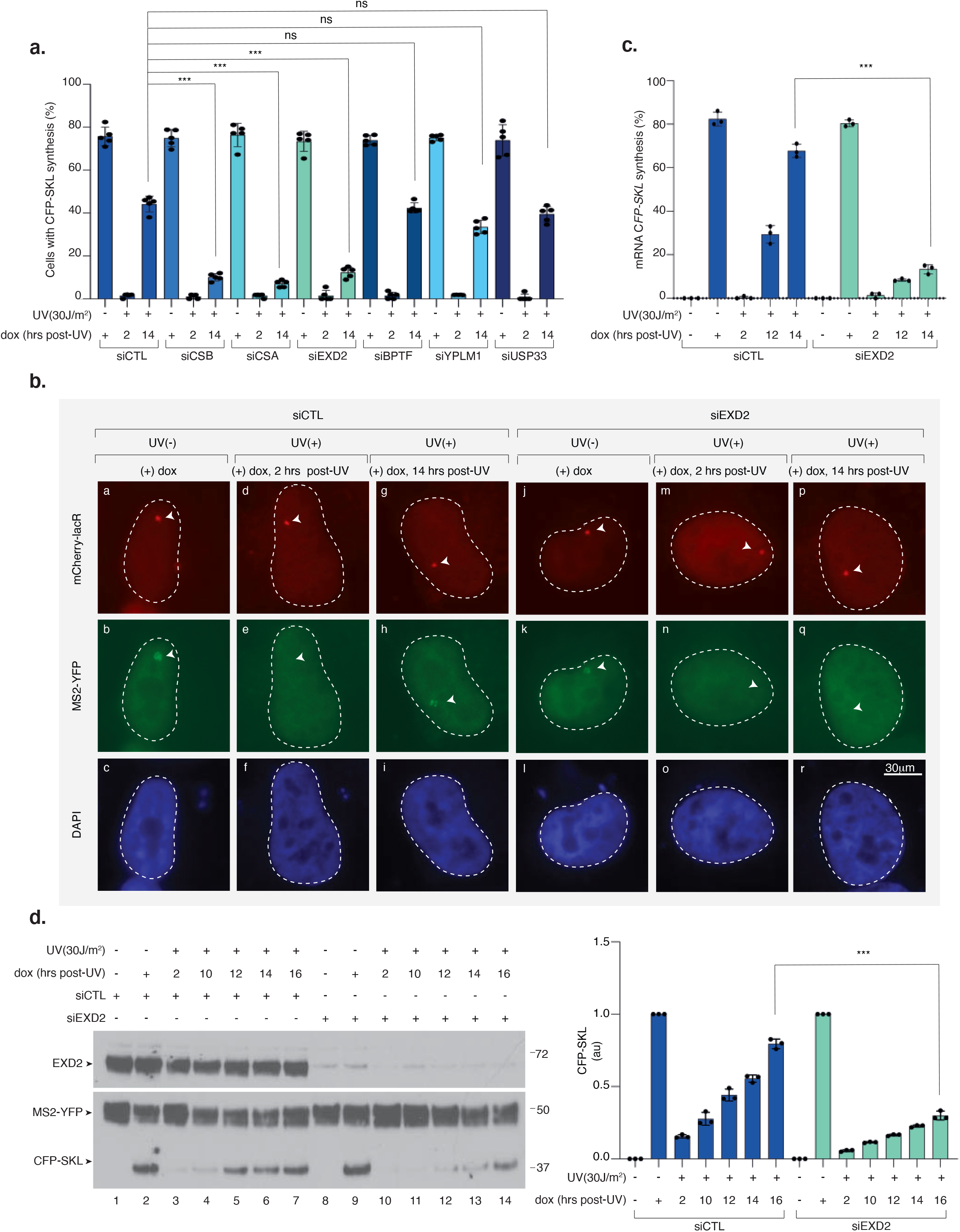
Knockdown of EXD2 impairs recovery of *CFP-SKL* expression after UV irradiation. a. Representative results from RNAi screen performed to identify new genes involved in transcription recovery after UV-irradiation. U-2 OS cells were transfected with siRNA for 24 hours, then with a construct expressing mCherry-lacR for 24 hours before UV irradiation (30J/m^2^) and subsequent 2-hour pulse-incubation with dox starting at various time points post-UV as indicated. Nascent *CFP-SKL* mRNAs were detected at the reporter locus by accumulation of the MS2-YFP protein to the MS2 RNA loop. Quantification of the transcribing locus was done and results are expressed as % of cells showing YFP-MS2 accumulation at a single locus (n= 250 cells in five independent experiments) (+/-SD). P-value (two-sided unpaired T-test) was extrapolated (***<0.005). b. Representative confocal images of cells treated with siCTL or siEXD2 as described above. Images of the cells were obtained with the same microscopy system and constant acquisition parameters. c. Cells were treated as described above and after the 2-hour pulse-incubation with dox, starting at various time points post-UV as indicated, the relative amount of *CFP-SKL* mRNA was quantified by RT-qPCR. Bars represent mean values of three different experiments (Biological triplicates) (+/-SEM). P-value (two-sided unpaired T-test) was extrapolated (***<0.005). d. U-2 OS cells were treated as described above. After the 2-hour pulse-incubation with dox, starting at various time points post-UV as indicated, the cells were let to recover for 4 hours before lysis. Extracts were resolved by SDS-PAGE and immuno-blotted with anti-GFP (recognizing both the MS2-YFP and CFP-SKL proteins) and anti-EXD2. Lanes 1 and 8 are negative controls in which cells were not treated with dox. Lanes 2 and 9 are positive controls in which cells were treated with dox for 2 hours before to recover 4 hours in the absence of dox. CFP-SKL signals were quantified using ImageJ software (NIH), normalized with YFP-MS2 signals and reported on the graph (1 is the value for dox (+) for siCTL or siEXD2). P-value (two-sided unpaired T-test) was extrapolated (***<0.005).

### EXD2 nuclease activity is required for RRS

To confirm the result of our screen, we next used HeLa cells in which EXD2 was stably knocked-down using shRNA (EXD2^-/-^-cl1) (Figure 2a, compare lanes 1 and 2). To measure global RRS, we pulse-labeled nascent mRNAs at various time points after UV irradiation (15J/m^2^) using 5-ethynyluridine (EU) (Alekseev et al., 2017). At this UV dose, all transcribed gene strands should contain at least one lesion that blocks RNAPII elongation (van Hoffen et al., 1995). We pre-treated the cells with a low concentration of actinomycin D (0.05µg/ml) to abolish the intense nucleolar EU staining due to RNAPI-dependent ribosomal RNA synthesis. In these conditions, EU incorporation mainly reflects RNAPII-dependent RNA transcription (Alekseev et al, 2017). Within the first hour after UV irradiation, we observed a strong inhibition (50%) of mRNA synthesis in both EXD2^+/+^ and EXD2^-/-^-cl1 cells (Figure 2b and Supplemental Figure 2a, panels a-b and e-f). In agreement with the above data, RRS was progressively restored in wild-type EXD2^+/+^ cells over time, whereas it remained deficient in EXD2^-/-^-cl1 cells (Figure 2b and Supplemental Figure 2a, compare panels c-d with g-h).

**Figure 2:**
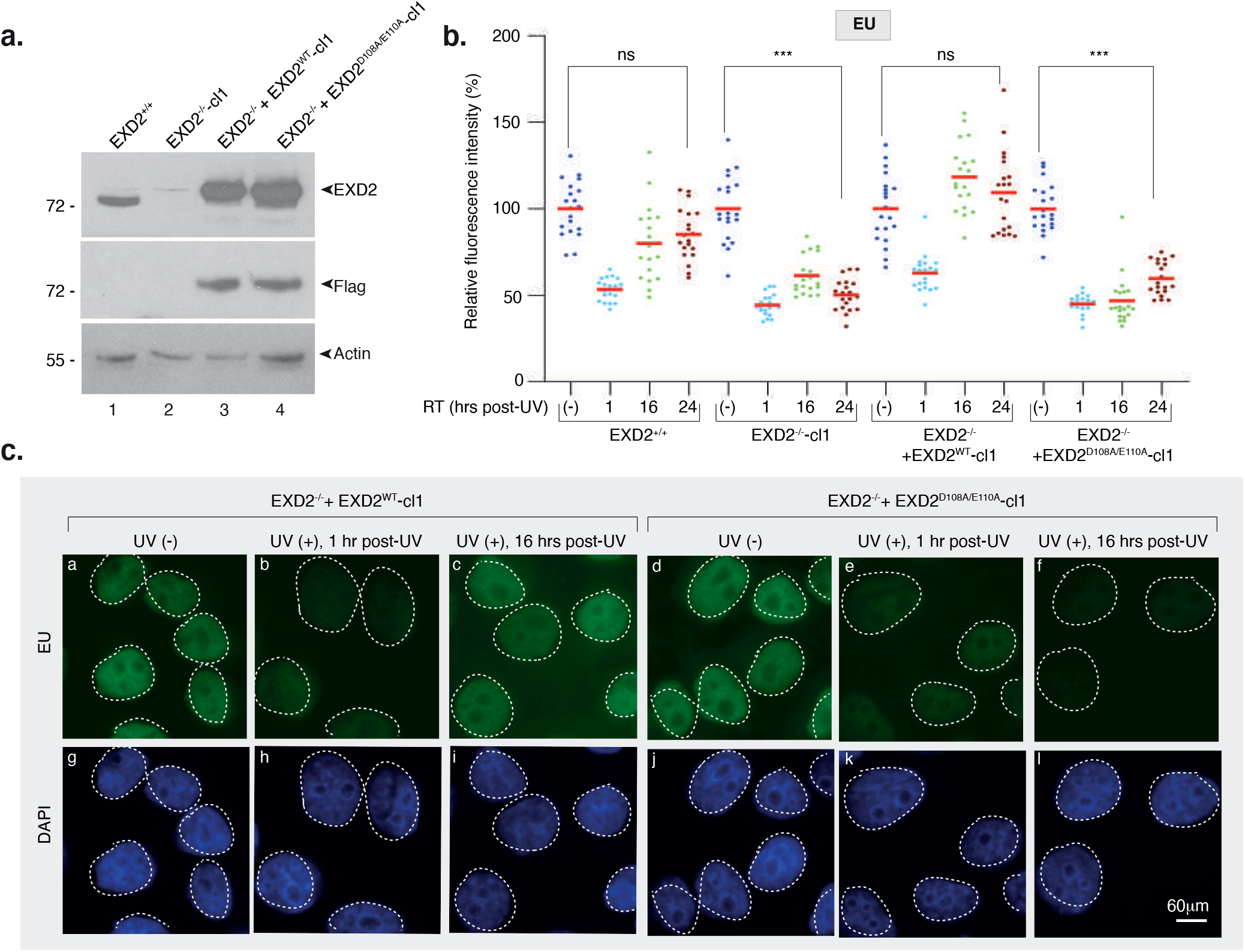
Lack of EXD2 nuclease activity inhibits RRS after UV irradiation. a. Protein lysates from EXD2^+/+^, EXD2^-/-^-cl1, EXD2^-/-^+EXD2^WT^-cl1 and EXD2^-/-^ +EXD2^D108A/E110A^-cl1 cells were immuno-blotted for proteins as indicated. Molecular mass of the proteins is indicated (kDa). b. Cells were mock- or UV-irradiated (15J/m^2^) and mRNA was labelled with EU at the indicated time points post-UV. EU signal was quantified by ImageJ and relative integrated densities, normalized to mock-treated level set to 100%, are reported on the graph (n=20 cells per conditions). Red bars indicate mean integrated density. P-value (two-sided unpaired T-test) was extrapolated (***<0.005). *RT*; recovery time. c. Representative confocal images of EXD2^-/-^+EXD2^WT^-cl1 and EXD2^-/-^+EXD2^D108A/E110A^-cl1 treated like in **(b)**. Images of the cells were obtained with the same microscopy system and constant acquisition parameters.

To explore the role of the exonuclease activity of EXD2 in RRS, EXD2^-/-^-cl1 cells were subsequently complemented with either shRNA-resistant FLAG-HA-tagged wild-type (EXD2^-/-^+EXD2^WT^-cl1) or dominant negative shRNA-resistant nuclease-dead EXD2 containing two substitutions at positions D108 and E110 (EXD2^-/-^+EXD2^D108A/E110A^-cl1). These two amino-acids are located in the active site of EXD2 and are known to be essential for its nuclease activity (Nieminuszczy et al., 2019), (Park et al., 2019). RRS was restored in EXD2^-/-^+EXD2^WT^-cl1 cells but not in nuclease-dead EXD2^-/-^+EXD2^D108A/E110A^-cl1 cells (Figure 2b and Figure 2c, compare panels a-c with d-f), showing that RRS requires the nuclease activity of EXD2. We noted that the stability of the RPB1 subunit of RNAPII after UV-irradiation was not affected by the depletion of EXD2 or the lack of its exonuclease activity (Supplemental Figure 2b). As noted above, we also observed that mRNA synthesis was indistinguishable in all four mock-treated HeLa cells, suggesting that EXD2 is not required for RNAPII-dependent transcription in the absence of genotoxic attack (Figure 2c, panels a and d, and Supplemental Figure 2a, panels a and e). Similar results were obtained with an additional set of HeLa clones (EXD2^-/-^-cl2, EXD2^-/-^+EXD2^WT^-cl2 and EXD2^-/-^+EXD2^D108A/E110A^-cl2) (Supplemental Figures 2c-d). Thus, the knockdown and overexpression studies complement one another to establish that the EXD2 exonuclease activity has a crucial function in RRS following UV irradiation.

In a second set of experiments, we evaluated the role of EXD2 in the response to various treatments provoking transcription arrest without generating DNA damage. We either treated the cells with the transcriptional inhibitor 5,6-dichloro-1-beta-D-ribofuranosylbenzimidazole (DRB) for 30 minutes (Alekseev et al., 2017) or incubated them for 15 minutes at 4°C to block transcription. Following the chase of DRB or the re-incubation at 37°C, we observed similar transcriptional recovery in EXD2^-/-^+EXD2^WT^-cl1 and EXD2^-/-^+EXD2^D108A/E110A^-cl1 (Supplemental Figure 3). Taken together, these results suggest that EXD2 specifically contributes to the global transcription recovery operating after a genotoxic stress such as UV irradiation.

### Lack of EXD2 nuclease activity leads to mild UV sensitivity

To further examine the consequences of a lack of EXD2 activity on cell homeostasis, we measured the UV sensitivity of EXD2^-/-^-cl1, EXD2^-/-^+EXD2^WT^-cl1 and EXD2^-/-^+EXD2^D108A/E110A^-cl1 cells in comparison with the parental EXD2^+/+^ cells as well as two UV-sensitive cell lines, namely the CS-B patient CS1ANSV cell line (in which the TC-NER factor CSB was deficient) (Kristensen et al., 2013) and the HeLa XPC^-/-^ (in which the GG-NER factor XPC was depleted) (Biard et al., 2005). Upon increasing doses of UV irradiation, knockdown of EXD2 activity resulted in hypersensitivity of EXD2^-/-^-cl1 and EXD2^-/-^+EXD2^D108A/E110A^-cl1 cells, compared to EXD^+/+^ and EXD2^-/-^+EXD2^WT^-cl1 cells (Figure 3a). Interestingly, UV sensitivity of EXD2^-/-^-cl1 and EXD2^-/-^+EXD2^D108A/E110A^-cl1 cells was similar to that found in the TC-NER deficient CS-B cells but not as pronounced as the one found in the highly sensitive GG-NER deficient XPC^-/-^ cells.

**Figure 3:**
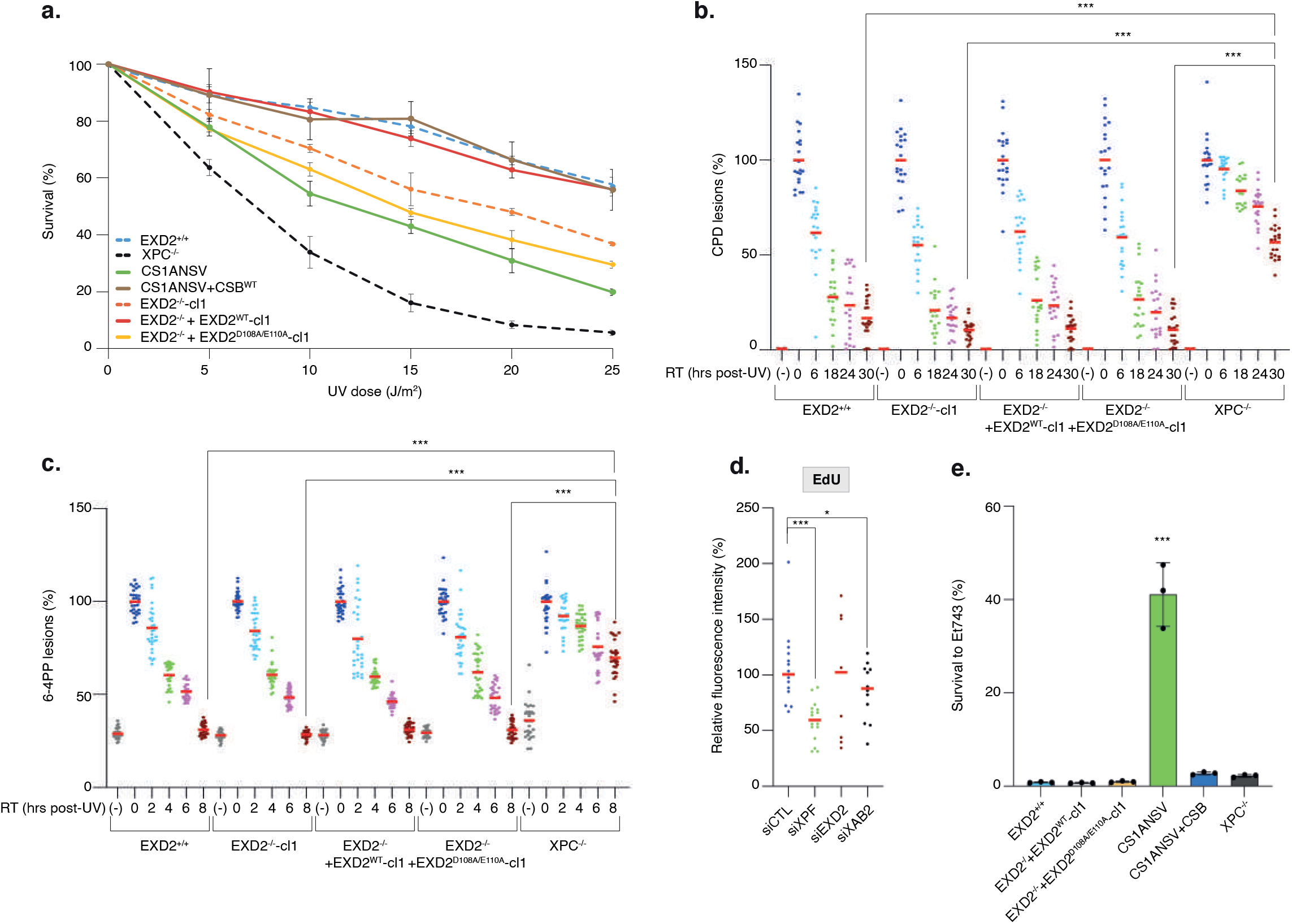
Lack of EXD2 nuclease activity sensitizes cells to UV irradiation. **a**. Cells were treated with increasing doses of UV irradiation and survival was determined 48 hours later. Data were normalized to the mock treatment controls (as value of 100). The values are the means of three independent experiments (+/-SEM) (Technical triplicates). **b-c**. Removal of UV lesions was measured in cells, harvested at different time points post-UV (15J/m^2^) as indicated. Cells were labelled with anti-CPD **(b)** or anti-6-4PP **(c)** antibodies and signals were quantified using ImageJ at the different times indicated in the figure. Graph represents the % of lesions remaining in the genome at different time points normalized to the amounts of lesions measured immediately after UV irradiation (as value of 100%). For each time point, a mean of 30 cells has been analysed. Red bars indicate mean integrated density. P-value (two-sided unpaired T-test) was extrapolated (***<0.005, ns=non-significant). RT; recovery time. (-); cells were mock-irradiated. 0; cells were irradiated and fixed immediately. d. GG-NER deficient XPC^-/-^ cells were treated either with siRNA against the TC-NER factor XAB2, the TC- and GG-NER factor XPF or against EXD2 for 24 hours before local UV irradiation (50J/m^2^) and EdU staining. The local TCR-UDS signals were quantified by ImageJ and reported on the graph. At least 15 cells were quantified for each situation. Red bars indicate mean integrated density. P-value (two-sided unpaired T-test) was extrapolated (***<0.005, *<0.01). e. Cells were treated with Et743 (0.5nM) and survival was determined 48 hours later. Data were normalized to the mock treatment controls (as value of 100). The values are the means of three independent experiments (+/-SEM) (Technical triplicates). ***P-value <0.005 (one-way ANOVA test).

To determine whether EXD2 was involved in the removal of UV-induced DNA damage by NER, we measured GG- and TC-NER in cells depleted of EXD2 activity. To this end, we first performed immunofluorescence-based quantification of UV lesions directly in cell nuclei (Alekseev et al., 2017). The removal rate of the two main types of UV lesions in EXD2^-/-^-cl1 and EXD2^-/-^+EXD2^D108A/E110A^-cl1 cells was higher to that of HeLa XPC^-/-^ cells and identical to that of EXD^+/+^ or EXD2^-/-^+EXD2^WT^-cl1 cells, implying that GG-NER was efficient in cells lacking EXD2 nuclease activity (Figures 3b-c). We used two different assays to measure TC-NER. We first performed unscheduled DNA repair synthesis (UDS) during TC-NER (TCR-UDS) (Donnio et al., 2019). Using GG-NER deficient XPC^-/-^ cells to ensure that repair replication in the UV-damage area was due to ongoing TC-NER, we measured repair replication via incorporation of EdU into newly synthesized DNA after local UV irradiation. Loss of the TC-NER specific factor XAB2 or TC/GG-NER factor XPF using siRNA induced similar deficiency in TCR-UDS, while loss of EXD2 had no impact (Figure 3d). We next used the particularity of TC-NER deficient cells to be resistant to the DNA binder and anti-cancer drug Ecteinascidin 743 (Et743) (Takebayashi et al., 2001). Indeed, the TC-NER deficient CS1ANSV cells showed high resistance to Et743 that was abolished in the recovered CS1ANSV+CSB cells (Figure 3e). In contrast, knockdown of EXD2 exonuclease activity did not impact the sensitivity of the corresponding cells to Et743. Finally, while γH2AX accumulated after UV-irradiation and persisted in NER deficient cells (Oh et al., 2011), no accumulation of γH2AX after knockdown of EXD2 was observed over time after UV irradiation (Supplemental Figure 4). Altogether, these results suggest that while the knockdown of EXD2 sensitizes cells to UV irradiation, the nuclease is not involved in GG-or TC-NER.

### EXD2 degrades nascent mRNA under synthesis at the time of UV irradiation

The above data point to a direct processing of mRNA by EXD2 nuclease activity during transcription recovery. To study this function, we first wanted to analyze the fate of mRNA under synthesis at the time of UV irradiation and developed the assay described in Figure 4a, upper panel. We inhibited ribosomal RNA synthesis with a low concentration of actinomycin D and subsequently labeled nascent mRNAs with a 10 minutes EU pulse. We then chased EU and immediately UV-irradiated the cells (15J/m^2^). Fixing them 1 hour or 16 hours post-chase, we were able to follow, during the recovery phase, the fate of mRNAs under synthesis when cells were subject to a genotoxic attack. In the four mock-treated cells, we observed a 50-40% reduced fraction size of EU-labeled mRNAs between 1 and 16 hours of culture (probably reflecting both the turn-over of mRNAs and their dilution during cell division) (Figure 4a and Supplemental Figure 5, compare panels a and c with panels i and k). Interestingly, UV irradiation of wild-type EXD2^+/+^ and EXD2^-/-^+EXD2^WT^-cl1 cells provoked a 70% reduced fraction size of EU-labeled mRNAs between 1 and 16 hours of culture, while EXD2^-/-^-cl1 and EXD2^-/-^+EXD2^D108A/E110A^-cl1 cells were refractory to this reduction and showed a situation similar to mock-treated cells with a 50-40% reduced fraction size of EU-labeled mRNAs (Figure 4a and Supplemental Figure 5, compare panels b and j with panels d and l).

**Figure 4:**
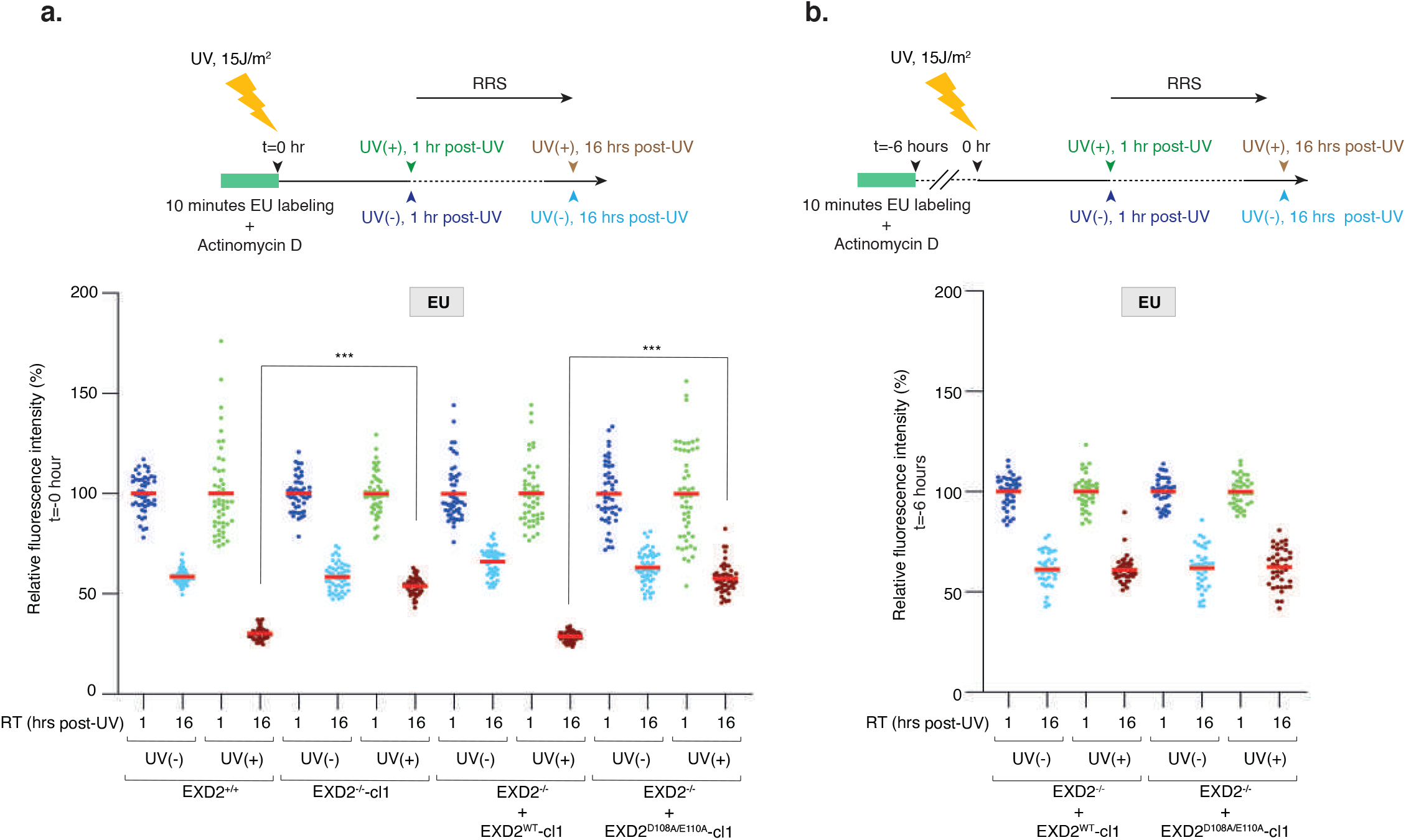
EXD2 degrades mRNA under synthesis at the time of UV irradiation. **a. Upper panel**; Scheme of the EU pulse-chase method used to analyze the fate of mRNA being synthesized at the moment of UV irradiation. Cells were incubated for 30 minutes with Actinomycin D (0.05µg/ml) to specifically inhibit RNAPI transcription and mRNAs were pulse-labeled with EU for 10 minutes prior to UV irradiation (15J/m^2^). Cells were let to recover for 1 hour or 16 hours post-UV before fixation. Actinomycin D was maintained during the experiment. **Lower panel;** Cells were treated as indicated in the upper panel and EU signals were quantified using ImageJ and normalized to the value obtained at 1 hour set to 100%. Values are reported on the graph (n=50 cells). P-value (two-paired T-test) was extrapolated (***<0.005). Red bars indicate mean integrated density. *RT*; recovery time. **b. Upper panel**; Compare to panel (a), UV irradiation (15J/m^2^) was performed 6 hours after EU labelling and cells were let to recover for either 1 hour or 16 hours post-UV before fixation. Actinomycin D was maintained during the experiment. **Lower panel;** Cells were treated as indicated in upper panel and EU signals were quantified using ImageJ. Values are reported on the graph (n=50 cells). P-value (two-paired T-test) was extrapolated and no statistically significant differences were detected at 16 hours, UV(+) between EXD2^-/-^ +EXD2^WT^-cl1 and EXD2^-/-^+EXD2^D108A/E110A^-cl1 cells. Red bars indicate mean integrated density. *RT*; recovery time.

In another set of experiments, we performed UV-irradiation long after EU labeling (6 hours) so that the labeled mRNAs were synthetized long before the UV treatment (Figure 4b, upper panel). In these conditions, the reduced fraction size of labelled mRNAs between 1 and 16 hours after irradiation was 50-40% for the four cell lines, regardless of whether EXD2 nuclease activity was present or not, a situation that resembles that of the mock treatment (Figure 4b). These experiments suggest that the EXD2 nuclease degrades, during the recovery phase, a large fraction of the nascent mRNAs (30%) that were being synthesized at the time of UV irradiation.

### UV irradiation induces a transient interaction between EXD2 and RNAPII

After having established the involvement of EXD2, during the recovery phase, in the degradation of mRNA under synthesis at the time of UV irradiation, we studied the potential connection of the nuclease with RNAPII. Using cell fractionation procedure, we first observed that the recruitment of EXD2 to chromatin increased in EXD2^-/-^+EXD2^WT^-cl1 cells upon UV irradiation (Figure 5a). Cell fractionation with or without pre-treatment with the transcriptional inhibitor DRB showed that EXD2 accumulated on chromatin even when transcription was inhibited, characterized by the loss of the phosphorylated and transcriptionally active form of RPB1 (RNAPIIO) (Figure 5b).

**Figure 5:**
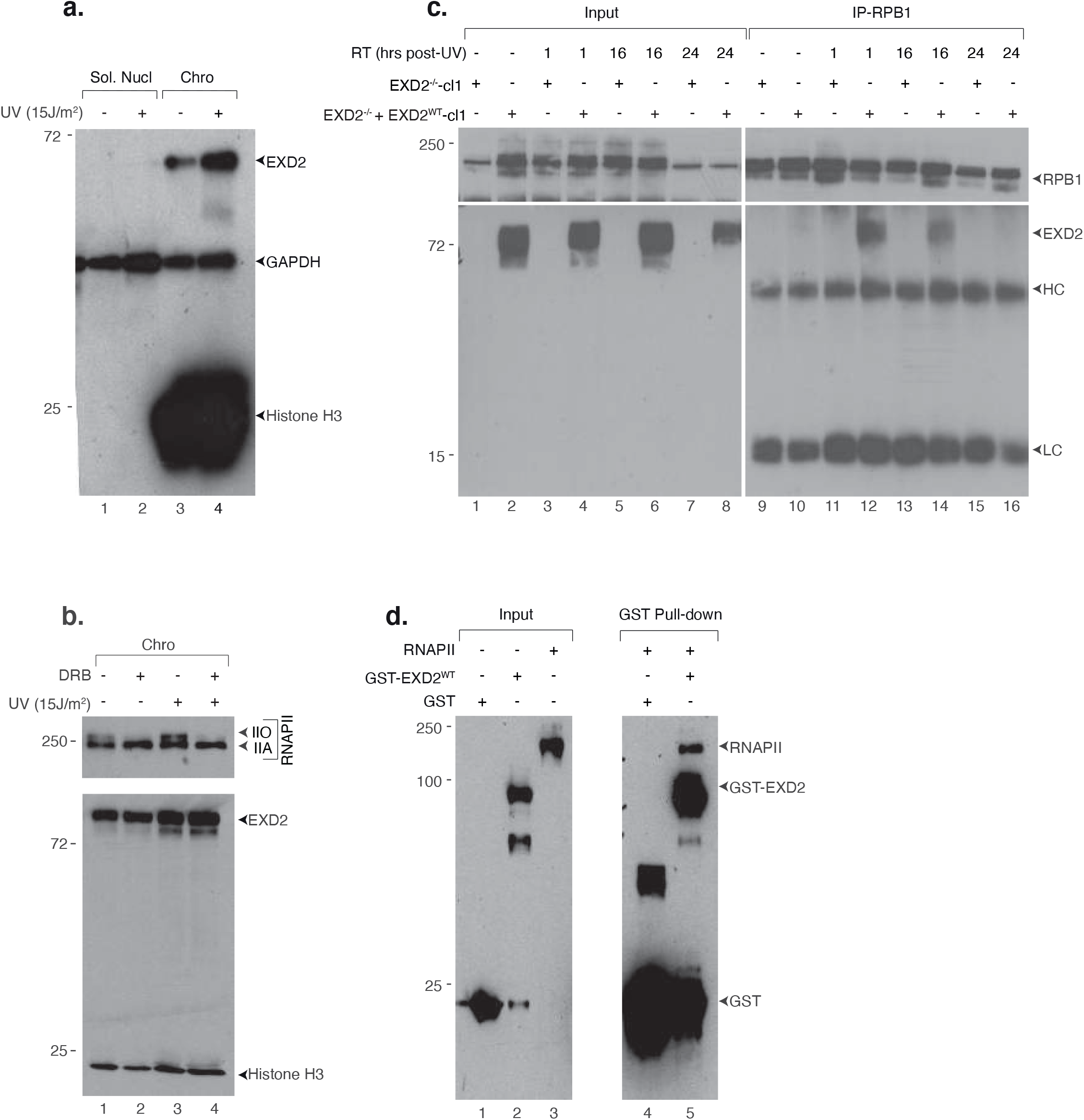
EXD2 transiently interacts with RNAPII after UV irradiation. a. EXD2^-/-^+EXD2^WT^-cl1 cells were mock- or UV-irradiated (15J/m^2^) and let to recover for an hour. Cells were fractionated in the soluble nuclear (Sol. Nucl) and chromatin (Chro) fractions, which were resolved by SDS-PAGE and immunoblotted against the indicated proteins. Molecular sizes are indicated (KDa). b. EXD2^-/-^+EXD2^WT^-cl1 cells were pre-treated or not with DRB (1 hour) before being mock- or UV-irradiated (15J/m^2^) and let to recover for an hour, post-UV. Cells were fractionated and the chromatin (Chro) fractions were resolved by SDS-PAGE and immunoblotted against the indicated proteins. Molecular sizes are indicated (KDa). Note that the Histone H3 bands appear different from panel **(a)** because different sources of anti-H3 were used in these two blots (**a**; anti-Histone H3 from mouse, **b**; anti-Histone H3 from rabbit). c. EXD2^-/-^-cl1 or EXD2^-/-^+EXD2^WT^-cl1 cells were mock- or UV-irradiated (15J/m^2^) and let to recover for the indicated period of time post-UV (RT). RNAPII was immunoprecipitated using anti-RPB1 from nuclear extracts and protein were resolved by SDS-PAGE and immunoblotted using anti-RPB1 or anti-EXD2 antibodies. *HC*, antibody heavy chain. *LC*, antibody light chain. *RT*; recovery time. d. Purified RNAPII from HeLa cells (Gerard et al., 1991) was incubated with recombinant pulldown GST-EXD2^WT^. Following washes, fractions were resolved by SDS-PAGE and immunoblotted against the indicated proteins. Controls IP was performed with GST alone (lane 4).

We next performed an immunoprecipitation of RNAPII in EXD2^-/-^+EXD2^WT^-cl1 cells before or after UV irradiation (15J/m^2^). We observed a transient kinetic of coprecipitation between EXD2 and the RPB1 subunit of RNAPII, which was clearly visible 1 hour post treatment (Figure 5c, lanes 9-12) and then begins to decrease at later time points to reach the level of mock-treated cells 24 hours after UV irradiation (compare lanes 12-14-16). We next expressed the full-length GST-tagged EXD2^WT^ in bacteria and performed a GST pull-down assay with purified RNAPII from HeLa cells (Gerard et al., 1991). GST-EXD2^WT^ pulldown co-precipitated RPB1, suggesting a direct interaction between EXD2 and RNAPII (Figure 5d). These data highlight a transient direct interaction between EXD2 and RNAPII taking place quickly after UV irradiation and persisting during the recovery phase.

### EXD2 interacts with a subset of RNAPII that stalls persistently on DNA

We then asked whether we could reconstitute EXD2 recruitment *in vitro* on an elongation-blocked RNAPII. We approached this question using an *in vitro* protein/DNA binding assay consisting of a biotinylated DNA template containing the AdMLP promoter and a transcribed region of 309 base pairs. The template was immobilized to streptavidin beads and incubated with purified RNAPII fraction from HeLa cells as well as with the recombinant general transcription factors (GTF: TFIIB, TBP, TFIIE, TFIIF, TFIIH) to form the pre-initiation complex (PIC). Bacterially purified recombinant EXD2 (rEXD2, without GST) was added at different stages of the assay (Figure 6a, left panel). Addition of NTP induced transcription initiation, whereas their subsequent chase induced RNAPII elongation arrest (Compe et al., 2019) (Figure 6a, middle panel). While western blot analysis of the remaining DNA-bound proteins revealed a very weak background signal of EXD2 to the DNA template in absence of RNAPII and its GTF (Figure 6a, right panel, lane 1), a clear recruitment of EXD2 occurred to the PIC in the absence of NTP (lane 3). In contrast, the presence of EXD2 did not improve the recruitment of RNAPII or GTFs (as observed for TFIIEα) (compare lane 2 and 3). The addition of NTP (lane 4) induced the initiation of transcription and the beginning of the elongation step characterized by the emergence of RNAPIIO as well as the release of the basal transcription factor TFIIEα from the DNA template (Figure 6a, middle panel) (Compe et al., 2019). Under these conditions of transcription elongation, EXD2 was released from the RNAPII complex (compare lane 3 with lane 5). Interestingly, the chase of NTP, which blocks RNAPII in elongation, caused EXD2 to be recruited again to the DNA template (compare lane 5 with lane 7). In another set of experiments, we determined whether NTP or ATP were required to induce EXD2 release from RNAPII during initiation. Addition of ATP triggered EXD2 release that was clearly enhanced by the presence of the four NTP (Figure 6b).

**Figure 6:**
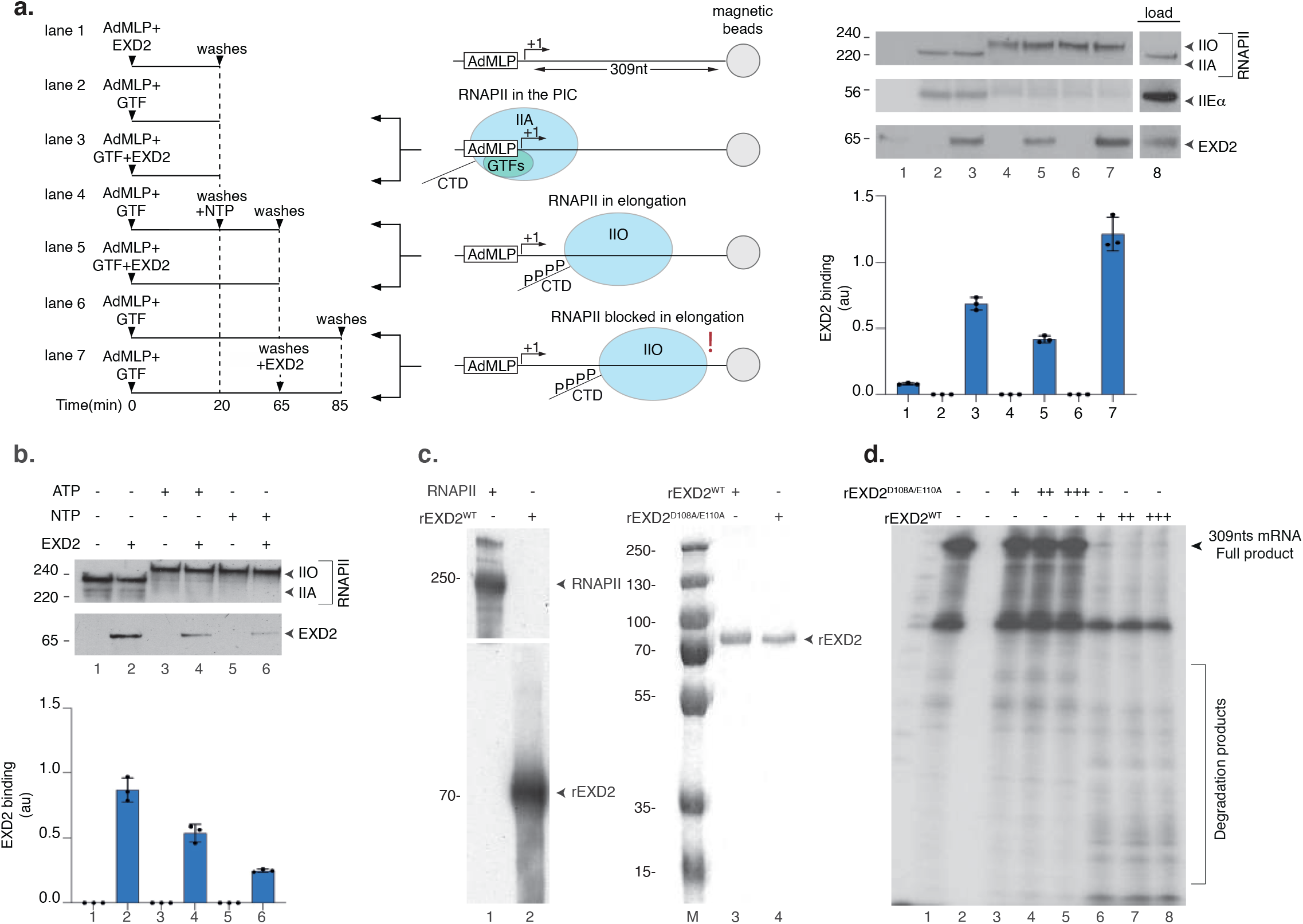
EXD2 preferentially interacts with RNAPII stopped in elongation. **a. Left Panel**; biotinylated DNA template, containing the AdML promoter upstream to a transcribed region of 309nts length, was bound to streptavidin magnetic beads and incubated for 20 minutes with purified RNAPII, TFIIB, TBP, TFIIF and TFIIH (GTF). After washes, NTPs were added to initiate RNAPII elongation for 45 minutes. Eventually, the NTPs were removed following additional washes and the reaction was allowed to continue for an additional 20 minutes. EXD2 was added at different times as indicated. **Middle panel**; the three conditions that were tested are summarized in the scheme; in the absence of NTPs where PIC is formed, in the presence of NTPs in which RNAPII is in elongation, and after the chase of NTPs in which RNAPII is blocked from elongating. **Right panel**; after the last washes, the binding of different factors was evaluated by immunoblotting. The loading proteins are indicated (load). The signals for EXD2 were quantified by Image J and plotted in arbitrary units (au). The values are the means of three independent experiments (+/-SD) (Technical triplicates). Molecular sizes are indicated (KDa). b. The biotinylated DNA template was bound to streptavidin magnetic beads and incubated for 20 minutes with purified RNAPII, TFIIB, TBP, TFIIF and TFIIH (GTF) with or without EXD2 as indicated. After washes, ATP or NTPs were added for 45 minutes. After washes, the binding of different factors was evaluated by immunoblotting and the signals for EXD2 were quantified by Image J and plotted in arbitrary units (au). The values are the means of three independent experiments (+/-SD) (Technical triplicates). Molecular sizes are indicated (KDa). **c. Left panel;** recombinant EXD2^WT^ purified from insect cells and RNAPII purified from HeLa were resolved by SDS-PAGE and immunoblotted with anti-EXD2 and anti-RPB1 antibodies. Molecular sizes are indicated (KDa). **Right panel;** Coomassie staining of recombinant EXD2^WT^ and EXD2^D108A/E110A^ purified from baculovirus infected cells. d. RNA (309nts) was transcribed from a linear template containing the AdML promoter using a reconstituted RNAPII run-off transcription assay containing recombinant TBP, TFIIB, TFIIE, TFIIF, TFIIH and purified RNAPII fraction from HeLa cells together with radiolabelled CTP for 30 minutes. Then, increasing amount of recombinant EXD2^WT^ or EXD2^D108A/E110A^ (10, 20 and 30 ng) was added to the reaction for an additional 10 minutes incubation period before the reaction was stopped.

We subsequently examined the impact of the exonuclease activity of EXD2 on newly synthesized mRNA. To detect mRNA, we complemented the above system with radio-labelled CTP (Alekseev et al., 2017). Note that the purified RNAPII fraction from HeLa cells was devoid of EXD2 (Figure 6c, left panel). Recombinant human EXD2^WT^ and nuclease-dead EXD2^D108A/E110A^ were purified from insect cells in parallel (Figure 6c, right panel). Transcription of the AdMLP-containing DNA template led to the production of an mRNA transcript of 309 nucleotides length in 30 minutes (Figure 6d, lane 1). Addition of increasing amounts of recombinant purified EXD2^WT^ for the last 10 minutes of the reaction induced the degradation of the mRNA transcript whereas EXD2^D108A/E110A^ had no impact (compare lanes 3-5 to 6-8). Taken together, these data suggest that a fraction of EXD2 is recruited to chromatin after UV-irradiation to directly interact with elongation-blocked RNAPII and degrade mRNA under synthesis.

## Discussion

Transcription is controlled in time and space by complex epigenetic and signaling-mediated regulatory networks at each step of the process. When cells are subjected to genotoxic attack, DNA damage impacts several crucial cellular functions, including transcription. Indeed, if these lesions are bulky and located in the transcribed strand of an active gene, they become a major complication during its transcription because they constitute a strong barrier to RNAPII forward translocation and result in its blockage, generating transcriptional genotoxic stress (Noe Gonzalez et al., 2021), (Jia et al., 2021). Cells cope with this stress firstly by inhibiting global gene expression, then by removing lesions that block RNAPII progression using the TC-NER pathway, and finally by initiating RRS at both promoters and damaged sites. How cells resume transcription after an acute genotoxic attack is crucial because inappropriate restarting is toxic and leads to cellular dysfunction and apoptosis, as observed in cells from patients with CS, which show intermediate sensitivity to UV irradiation coupled with a defect in RRS (Gregersen and Svejstrup, 2018). With this in mind, we sought to find genes involved in RNAPII-dependent gene expression recovery after genotoxic attack and unveiled a key role for the 3’-5’ exonuclease activity of EXD2 in this process. Recent studies have shown that RNAPI-dependent ribosomal gene transcription is also blocked shortly after a genotoxic stress and recovers over time. A TC-NER machinery removes lesions in ribosomal genes with the participation of CSA and CSB (Daniel et al., 2018). We tested whether EXD2 was involved in RNAPI-dependent transcription recovery but did not detect a defect in this process in cells lacking EXD2 (Dr. Mari-Giglia, personal information), suggesting a specific involvement of EXD2 in RNAPII-dependent transcription recovery after UV irradiation.

Cells lacking EXD2 nuclease activity exhibited intermediate sensitivity to UV irradiation, reminiscent of the phenotype observed in TC-NER deficient cells (Jia et al., 2021), (van den Heuvel et al., 2021). At first glance, this could indicate that EXD2 is involved in removing DNA lesions that block RNAPII during elongation. However, sensitivity to Et743, which required an active TC-NER pathway (Takebayashi et al., 2001), and the TCR-UDS assay indicate that TC-NER is unaffected by the absence of EXD2, suggesting an uncoupling of efficient TC-NER from deficient RRS in these conditions. Recently, regulation of the RNAPII pool by ubiquitination was shown to be central in the inhibition and restart of transcription in response to genotoxic stress (Tufegdžić Vidaković et al., 2020), (Nakazawa et al., 2020). Persistent depletion of RNAPII was shown to be largely responsible for the lack of transcriptional recovery observed in CS-B cells. Under our conditions, the RNAPII pool was not affected after UV irradiation by the lack of EXD2 nuclease activity, and we ruled out that a direct impact of EXD2 on RNAPII stability was involved in the RRS defect observed in EXD2-deficient cells. Instead, our observations suggest a direct action of EXD2 nuclease activity on mRNA, which was observed both *in vivo*, using pulse-labeling of nascent mRNA, and *in vitro*, using the transcription/nuclease run-off assay. These assays clearly show that the EXD2 nuclease processes, during the recovery phase, a fraction of mRNA representing 30% of the nascent mRNA being synthesized at the time of the genotoxic attack. During RNAPII backtracking, the ability of RNAPII to cleave its transcript potentially allows transcription to resume and cells to survive when the lesions are removed. A plethora of transcription factors, such as TFIIS, are known to stimulate transcript cleavage (Sigurdsson et al., 2010). Similarly, it seems reasonable to suggest that the exonuclease activity of EXD2 is involved in the mRNA processing associated with RNAPII backtracking in front of DNA lesion as illustrated by the dynamic interaction we observed between them, which transiently kicks in after genotoxic attack. Moreover, this activity is likely limited to a genotoxic attack situation because mRNA transcription was efficiently recovered after DRB or cold-shock treatment in cells lacking EXD2. Our observations also suggest that EXD2 is not involved in transcription *per se*, as EU incorporation or reporter expression was normal in mock-treated cells and cell viability was not affected by EXD2 knockdown in the absence of genotoxic stress (our data and (Nieminuszczy et al., 2019)). This is similar to other genes encoding transcript cleavage stimulatory factors, such a TFIIS, that are not essential for cell viability in the absence of genotoxic stress (Sigurdsson et al., 2010), which may reflect a potential redundancy in the function of these factors.

RNAPII backtracking in front of a lesion likely occurs over several nucleotides, such that the 3’ end of the RNA is no longer aligned with the RNAPII active site, preventing transcription restart. Therefore, the data presented here advocate for a scenario in which EXD2 transiently associates with an RNAPII that is stopped persistently on a gene during elongation by the presence of a transcription-blocking lesion, to potentially assist RNAPII in degrading mRNA from 3’ to 5’ when backtracking occurs (Geijer and Marteijn, 2018), (Noe Gonzalez et al., 2021). This activity, alone or in combination with that of RNAPII, could reactivate backtracked RNAPII by providing a new 3’ end to the mRNA to realign RNAPII active site with the ongoing mRNA. Why cells would require the 3’ to 5’ exonuclease activities of RNAPII and EXD2 to process mRNA at a damaged site is unclear, but consistent with this scenario, EXD2 is essential for cell viability after UV irradiation but is not required for NER to occur, demonstrating that UV sensitivity reflects the toxicity of the absence of RRS rather than a defect in DNA damage removal. To better understand the molecular mechanism of EXD2 involvement in RRS, the association of EXD2 with RNAPII was reconstituted *in vitro* using a transcribed DNA template and highly purified and recombinant transcription factors. Consistent with our hypothesis, we observed that elongation-active RNAPII associated less with EXD2 than elongation-blocked RNAPII on transcribed DNA. It is known that EXD2 discriminates RNA and DNA substrates via metal coordination (Mn2+ *vs* Mg2+) (Park et al., 2019). We show that under the physicochemical conditions allowing *in vitro* transcription and the presence of Mg2+, EXD2 exonuclease activity processes newly synthesized long mRNA molecules (309nts length), reinforcing our model of UV-induced recruitment of EXD2 to stalled RNAPII, followed by degradation of nascent mRNA before transcription resumes. Interestingly, these data are also the only ones to assign a ribonuclease function to EXD2 in the nucleus since previous work implicated it in the degradation of nuclear DNA either during DNA double strand break resection in non-homologous end joining or in the protection of stressed replication forks (Broderick et al., 2016), (Biehs et al., 2017), (Nieminuszczy et al., 2019).

## Acknowledgements

We thank Dr. Patrick Reilly for critical reading of the article, Dr. Wojciech Niedzwiedz (The institute of Cancer Research) for HeLa clones and recombinant EXD2 proteins and Dr. Roger Greenberg (University of Pennsylvania) for the U-2 OS-pTuner263 cells. This study was supported by ANR (TFIIH-2021), by the Ligue contre le cancer (Equipe Labélisée 2022-2024) and the grant ANR-10-LABX-0030-INRT, a French State fund managed by the Agence Nationale de la Recherche under the frame program Investissements d’Avenir ANR-10-IDEX-0002-02. M.C is supported by the “Ligue contre le Cancer”. C.E by the “Région Réunion”. LMD is supported by Agence Nationale de la Recherche (ANR-14-CE10-0009) and Institut National du Cancer (PLBIO17-043 and PLBIO19-126).

## Author contribution

J.S, M.C, P.C, L-M D, C.E, C.B and S.A conducted the experiments. J.S, JM-E, G M-G and E.C analyzed the results. F.C designed the experiments and wrote the paper.

## Material and Methods

### Cell culture

U-2 OS cells were cultured in DMEM (1g/l Glucose) containing 10% FCS and gentamycin. U-2 OS pTuner 263 cells were cultured in DMEM (1g/l Glucose) containing 10% Tet-system approved FBS, 1% Penicillin/Streptomycin, 400 µg/ml G418, and 100 µg/ml hygromycin B and 2µg/ml puromycin. The clones of each cell type (HeLa EXD2^-/-^-cl1 and 2, HeLa EXD2^-/-^+EXD2^WT^-cl1 and 2 and HeLa EXD2^-/-^+EXD2^D108A/E110A^-cl1 and 2) were cultured in DMEM (1g/l Glucose) containing 10% FCS and gentamycin supplemented with 0.25µg/ml of puromycin for HeLa EXD2^-/-^+EXD2^WT-^-cl1 and 2 and HeLa EXD2^-/-^+EXD2^D108A/E110A-^-cl1 and 2. XP4PA-SV, CS1ANSV, CS1ANSV+CSB and XPC Hela Silencix were cultured as described (Donnio et al., 2019), (Le May et al., 2010), (Kristensen et al., 2013).

### Quantification of actively transcribing cells

U-2 OS pTuner 263 cells were induced by 1µg/ml of dox for the indicated time intervals. Cells were fixed and the number of cells harboring an YFP-MS2 spot counted.

### *CFP-SKL* mRNA quantification

Total RNA was purified using TriReagent following the manufacturer’s protocol (Molecular Research Center, TR118) and cDNA was prepared by the SuperScript IV kit (Invitrogen, 18090050). qPCR reactions were carried out using the LightCycler480 (Roche) machine and the LightCycler 480 SYBR Green I Master (Roche, 04887352001).

### EU incorporation assay/RRS assay

RNA labeling by EU incorporation was performed with Click-iT RNA Alexa Fluor 488 Imaging Kit (Invitrogen, C10329) following the manufacturer protocol with the following modifications; 5EU was used at 0.1mM and labeling was performed during 1 hour to obtain a good linear EU signal as a function of the incubation time (Alekseev et al., 2017). Microscopy pictures were taken with Leica DM 4000 B equipped with a CoolSnap FX monochrome camera and EU signal intensity was quantified by ImageJ software.

### Immunofluorescence-based DNA lesion quantification

Cells were plated in a 24-well plate. 24 hours later, cells were UV-irradiated with UV-C lamp (15J/m^2^) and recovered for different recovery time intervals at 37°C, 5% CO2. Immuno-labeling of cyclobutane pyrimidine dimers (CPD) and 6-4 photoproducts (6-4PP) was performed using mouse anti-CPD and anti-6-4PP antibodies. DNA was denatured with 2M HCl for 20 minutes at RT and blocked in 10% FCS in PBS for 30 minutes prior to labeling. Microscopy pictures were taken with Leica DM 4000 B equipped with a CoolSnap FX monochrome camera and EU signal intensity was quantified by ImageJ software to determine the percentage of CPD and 6-4PP removal (100% represents the % of lesions measured just after UV irradiation).

### Transfections

Plasmid transfections were conducted using X-tremeGene DNA Transfection Reagent (Roche) according to manufacturer’s protocols. siRNA transfections were conducted using Lipofectamine RNAiMAX Transfection Reagent (Invitrogen) according to manufacturer’s protocols.

### TCR-Unscheduled DNA synthesis (TCR-UDS)

GG-NER-deficient XP4PA-SV cells (XP-C) were grown on 18 mm coverslips. siRNA transfections were performed 24 hours and 48 hours before TCR-UDS assays. After local irradiation at 50J/m^2^ with UV-C through a 5µm pore polycarbonate membrane filter, cells were incubated for 8 hours with 5-ethynyl-2’- deoxyuridine (EdU), fixed and permeabilized with PBS and 0.5% triton X-100. Then, cells were blocked with PBS+ solution (PBS containing 0.15% glycine and 0.5% bovine serum albumin) for 30 minutes and subsequently incubated for 1 hour with mouse monoclonal anti-γH2AX antibody 1:500 diluted in PBS. After extensive washes with PBS containing 0.5% Triton X100, cells were incubated for 45 minutes with secondary antibodies conjugated with Alexa Fluor 594 fluorescent dyes (Molecular Probes, 1:400 dilution in PBS). Next, cells were washed several times and then incubated for 30 minutes with the Click-iT reaction cocktail containing Alexa Fluor Azide 488. After washing, the coverslips were mounted with Vectashield containing DAPI (Vector). Images of the cells were obtained with the same microscopy system and constant acquisition parameters. Images were analyzed as follows using ImageJ and a circle of constant size for all images: (i) the background signal was estimated in the nucleus (avoiding the damage, nucleoli and other non-specific signal) and subtracted, (ii) the locally damaged area was defined by using the yH2AX staining, (iii) the mean fluorescence correlated to the EdU incorporation was then measured and thus an estimate of DNA synthesis after repair was obtained. For each sample three independent experiments were performed.

### Cell sub-fractionation

Cell lysis was performed using triton X-100 on either mock or UV-irradiated cells (10J/m^2^). Cells were subsequently trypsinised 1 hour after irradiation and washed twice with PBS. The following procedures were carried out at 4°C. The soluble fraction was collected by treatment with 5× volume 0.1% Triton X-100, 130mM KCl, 10mM K2 PO4 (pH 7.4), 10mM Na2 HPO4 (pH 7.4), 1mM Na_2_ATP (Grade II Sigma), 2.5mM MgCl2, 10% glycerol, 0.5mM DTT for 10 minutes followed by centrifugation at 4K RPM for 5 minutes. The remaining chromatin fraction was resuspended in 5x volumes of 2x Laemli-SDS-sample buffer and homogenized by sonification.

### Protein/DNA binding assay

Biotinylated AdMLP DNA template bound to streptavidin magnetic beads was incubated 20 minutes at 25°C with purified RNAPII, TFIIA, IIB, IIF, TBP, IIH and EXD2 in transcription buffer (20mM HEPES (pH 7.9), 7mM MgCl_2_, 55mM KCl). After three washings at 50mM NaCl, bound fractions were resolved by SDS-PAGE for immunoblottings and others were incubated 45 minutes at 25°C with NTP (200µM). After washings, these fractions were in turn resolved by SDS-PAGE or were further incubated 20 minutes with EXD2. The abundance of EXD2 was assessed by immunoblot densitometry analysis (using ImageJ software). Each signal was quantified three times and plotted in arbitrary units (au).

### Reconstituted runoff transcription

Reaction mixtures of 12μL containing 50ng of linear AdMLP DNA template and recombinant TFIIH, TFIIB, TFIIE, TFIIF, TBP together with purified RNA pol II as described (Coin et al., 2006) were pre-incubated for 20 minutes at 25°C in transcription buffer (20mM HEPES (pH7.9), 7mM MgCl_2_, 55mM KCl) and transcription was initiated by the addition of 2μL nucleotide solution to final concentrations of 600μM UTP, ATP, GTP and 0.6μM [α-^32^P] CTP. Reactions were carried out for 30 minutes and recombinant EXD2 was added for another 10 minutes. Reaction was stopped by the addition of 0.5μL of 0.5M EDTA (pH 8). The resulting RNA transcripts were analyzed on an 8% denaturing polyacrylamide gel.

### Statistics and reproducibility

Experimental data was plotted and analyzed using GraphPad Prism (GraphPad Software Inc.). The number of samples and replicates are indicated in the respective figure legends.

### Lead contact

Further information and requests for resources and reagents should be directed to and will be fulfilled by the Lead Contact, Dr. Frédéric Coin.

---

**Supplemental Figure 1:**
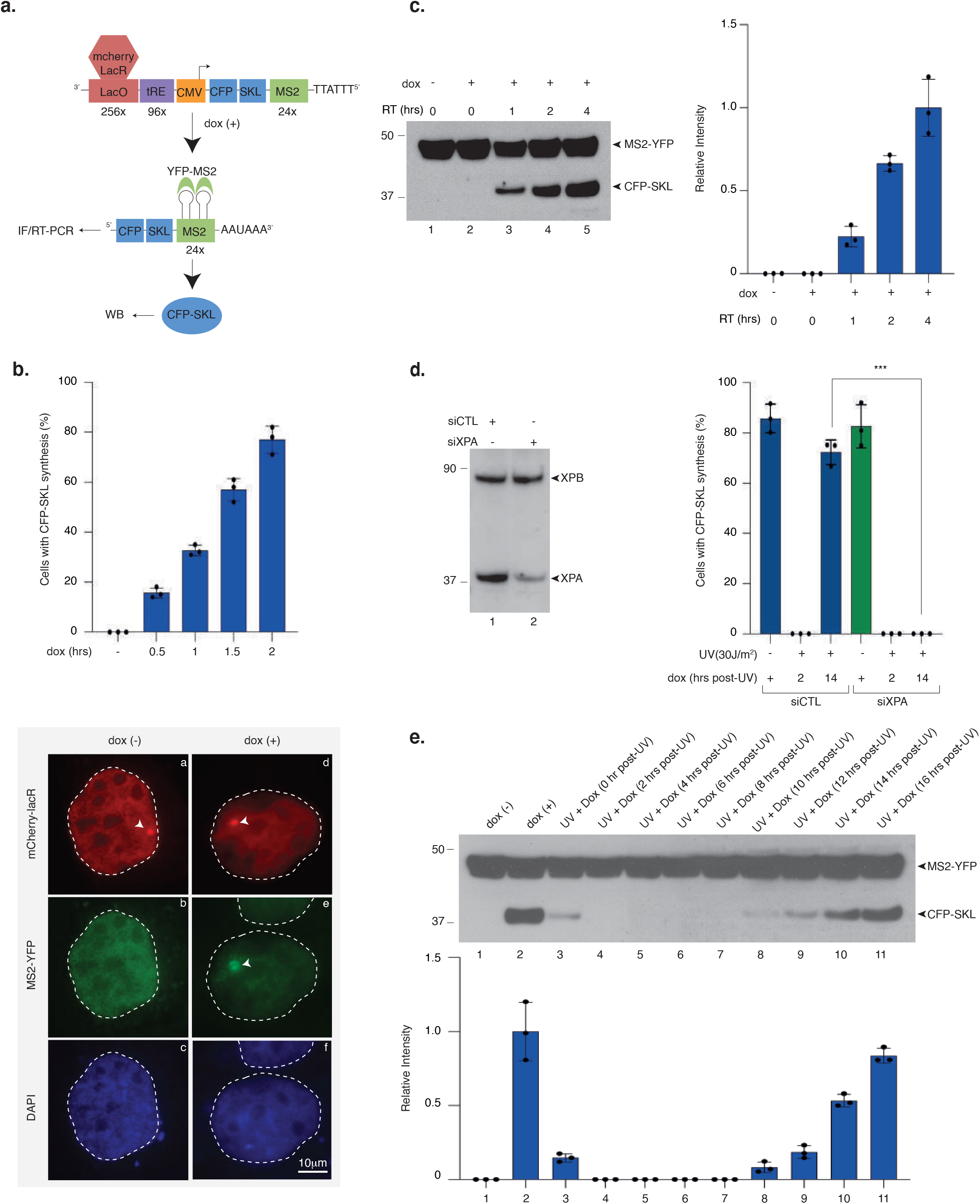
The CFP-SKL reporter assay. a. Scheme of the reporter locus. Briefly, 256 copies of the lactose operator (lacO) array provide examination of the reporter through a fluorescent mCherry protein fused to the lactose repressor (mCherry-lacR). Between LacO and the minimal CMV promoter, tandem repeats of tetracycline response elements (TREs) controlled the expression of the unique *CFP-SKL* mRNA transcript upon doxycycline (dox) treatment. The *CFP-SKL* mRNA sequence contained 24 repeats of the MS2 stem loop structure to monitor transcription through binding of the yellow fluorescent protein tagged MS2 viral coat protein (YFP-MS2) to the nascent transcript. The mRNA encoded a CFP-tagged peroxisomal targeting peptide (CFP-SKL) that can be detected by WB using an anti-GFP antibody. b. U-2 OS were treated with dox (1μg/ml) for different times as indicated and *CFP-SKL* mRNA expression was quantified using MS2-YFP accumulation (upper panel). Results are expressed as % of cells showing YFP-MS2 accumulation at a single locus (n= 3×20 cells, biological triplicates) (+/-SD). Representative confocal images of MS2-YFP accumulation at newly synthetized *CFP-SKL* mRNA, after 2-hour dox treatment, is shown (lower panel). The reporter locus can be detected with a mcherry-LacR fusion construct transfected for 24 hours before dox treatment, which recognized the LacO sequence. c. Immuno-blot for CFP-SKL and YFP-MS2 in U-2 OS cells treated with dox for 2 hours and subsequently let to recover (Recovery Time, RT) in dox-free medium for the indicated period of time. Following SDS-PAGE, extracts were immuno-blotted with anti-GFP. The OD of the CFP-SKL signals were quantified using ImageJ software (NIH), normalized with YFP-MS2 signals (+/-SEM). Three independent immunoblots were performed (technical triplicates). d. U-2 OS cells were transfected with siRNA for 24 hours, then with a construct expressing mCherry-lacR for 24 hours before UV irradiation (30J/m^2^) and subsequent 2-hour pulse-incubation with dox starting at various time points post-UV as indicated. Newly transcribed *CFP-SKL* mRNAs were detected at the reporter locus by accumulation of the MS2-YFP protein to the MS2 RNA loop. Quantification of the transcribing locus was done and results are expressed as % of cells showing YFP-MS2 accumulation at a single locus (n= 3×20 cells, biological triplicates) (+/-SD). e. U-2 OS cells were UV-irradiated (30 J/m^2^) and subsequent pulse-incubated for 2 hours with dox starting at various time points post-UV as indicated in brackets. Cells were allowed to recover for 4 hours in the absence of dox. Following SDS-PAGE, extracts were immuno-blotted with anti-GFP antibody. Lane 1 is a negative control in which cells were not treated with dox. Lane 2 is a positive control in which cells were treated with dox for 2 hours before to recover 4 hours in the absence of dox. The OD of the CFP-SKL signals were quantified using ImageJ software (NIH), normalized with YFP-MS2 signals and reported on the graph (+/-SEM). Three independent immunoblots were performed (technical triplicates).

**Supplemental Figure 2:**
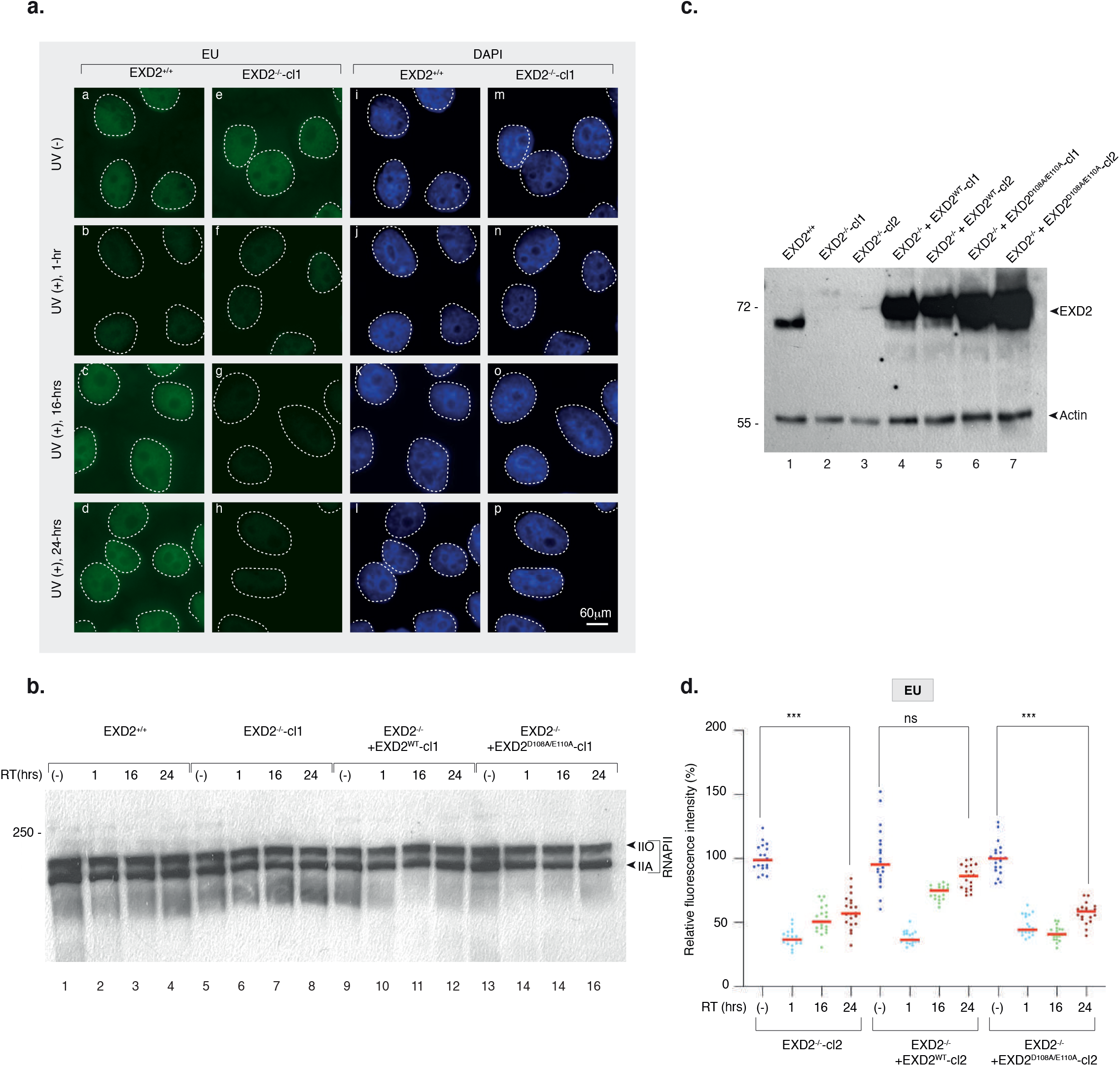
Lack of EXD2 induces inhibition of RRS after UV irradiation. a. Representative confocal images of EXD2^+/+^ and EXD2^-/-^-cl1 showing mRNA synthesis in different conditions. Cells were mock or UV-irradiated (15J/m^2^) and mRNA was labelled with EU 1, 16 and 24 hours after UV-irradiation. Images of the cells were obtained with the same microscopy system and constant acquisition parameters. b. Cells were mock or UV-irradiated (15J/m^2^) and let to recover for the indicated times before lysis. Protein lysates were immuno-blotted for RPB1 subunit of RNAPII. Molecular mass of the proteins is indicated on the left (kDa). RT; recovery time. IIO, phosphorylated form of RPB1. IIA non-phosphorylated form. c. Protein lysates either from EXD2^+/+^, EXD2^-/-^-cl1, EXD2^-/-^-cl2, EXD2^-/-^ + EXD2^WT^-cl1, EXD2^-/-^ + EXD2^WT^-cl2, EXD2^-/-^ + EXD2^D108A/E110A^-cl1 and EXD2^-/-^ + EXD2^D108A/E110A^-cl2 were immuno- blotted for proteins as indicated. Molecular mass of the proteins is indicated (kDa). d. Cells were mock or UV-irradiated (15J/m^2^) and mRNA was labelled with EU 1, 16 and 24 hours after UV-irradiation. EU signal was quantified by ImageJ and relative integrated density normalized to mock-treated level set to 100% are reported on the graph (n=20 cells per conditions). Red bars indicate median integrated density. P-value (two-sided unpaired T-test) was extrapolated (***<0.005).

**Supplemental Figure 3:**
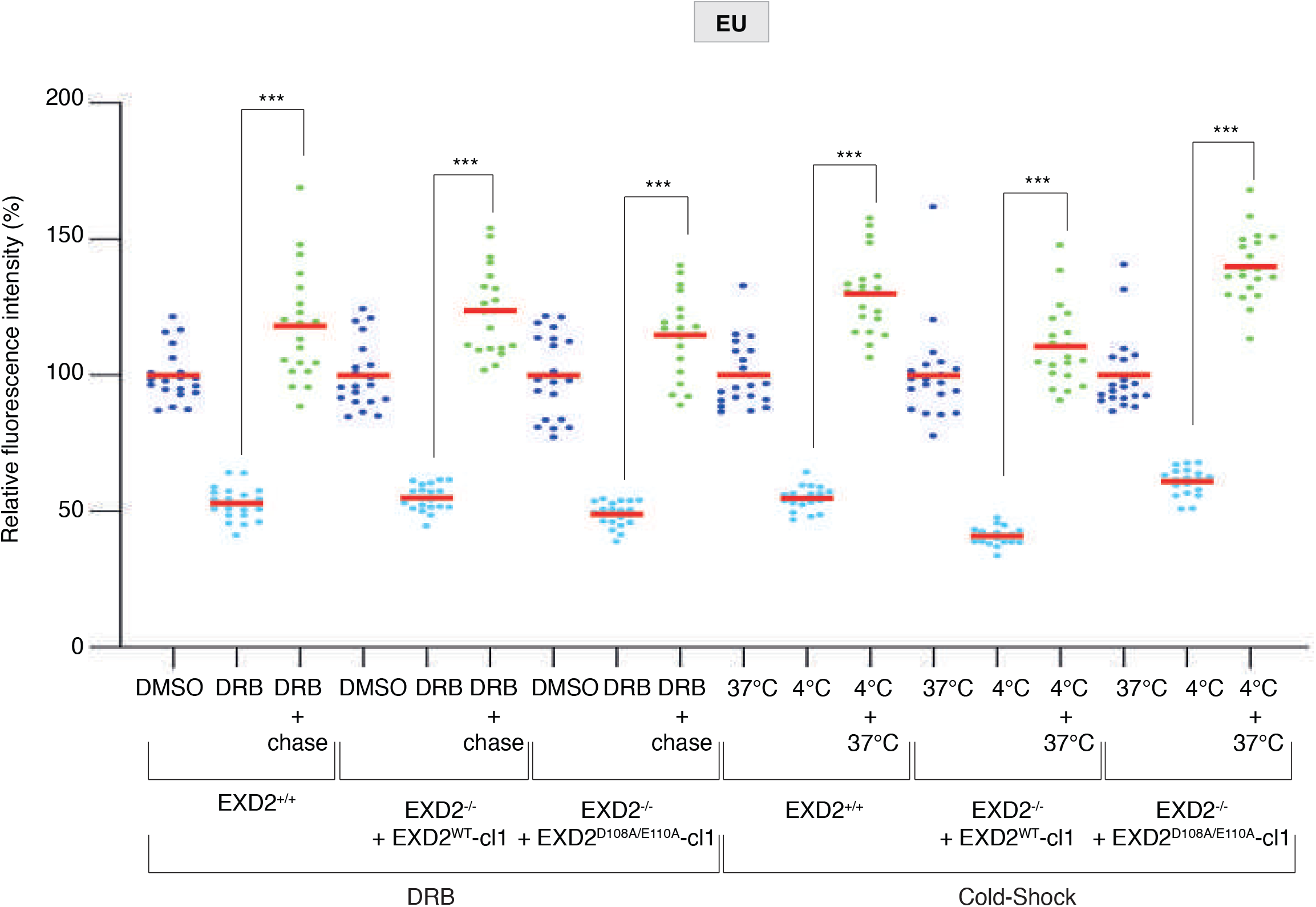
EXD2 is not involved is RRS after DRB or cold-shock treatment. **DRB:** EXD2^+/+^, EXD2^-/-^ + EXD2^WT^-cl1 and EXD2^-/-^ + EXD2^D108A/E110A^-cl1 were treated with DRB (100 µM) for 1 hour before chase and cells were let to recover for 30 minutes. mRNA was labelled with EU before (DMSO), during (DRB) or after the chase (DRB + chase) of DRB treatment. **Cold-Shock**: EXD2^+/+^, EXD2^-/-^ + EXD2^WT^-cl1 and EXD2^-/-^ + EXD2^D108A/E110A^-cl1 were incubated at 4°C for 15 minutes and let to recover at 37°C for 30 minutes. mRNA was labelled with EU before (37°C), after 15 minutes at 4°C (4°C) or after the recovery at 37°C (4°C+37°C). For both experiments, EU signal was quantified by ImageJ and relative integrated densities normalized to mock-treated level set to 100% are reported on the graph (n=20 cells per conditions). Red bars indicate average integrated density. P-value (two-sided unpaired T-test) was extrapolated (***<0.005).

**Supplemental Figure 4:**
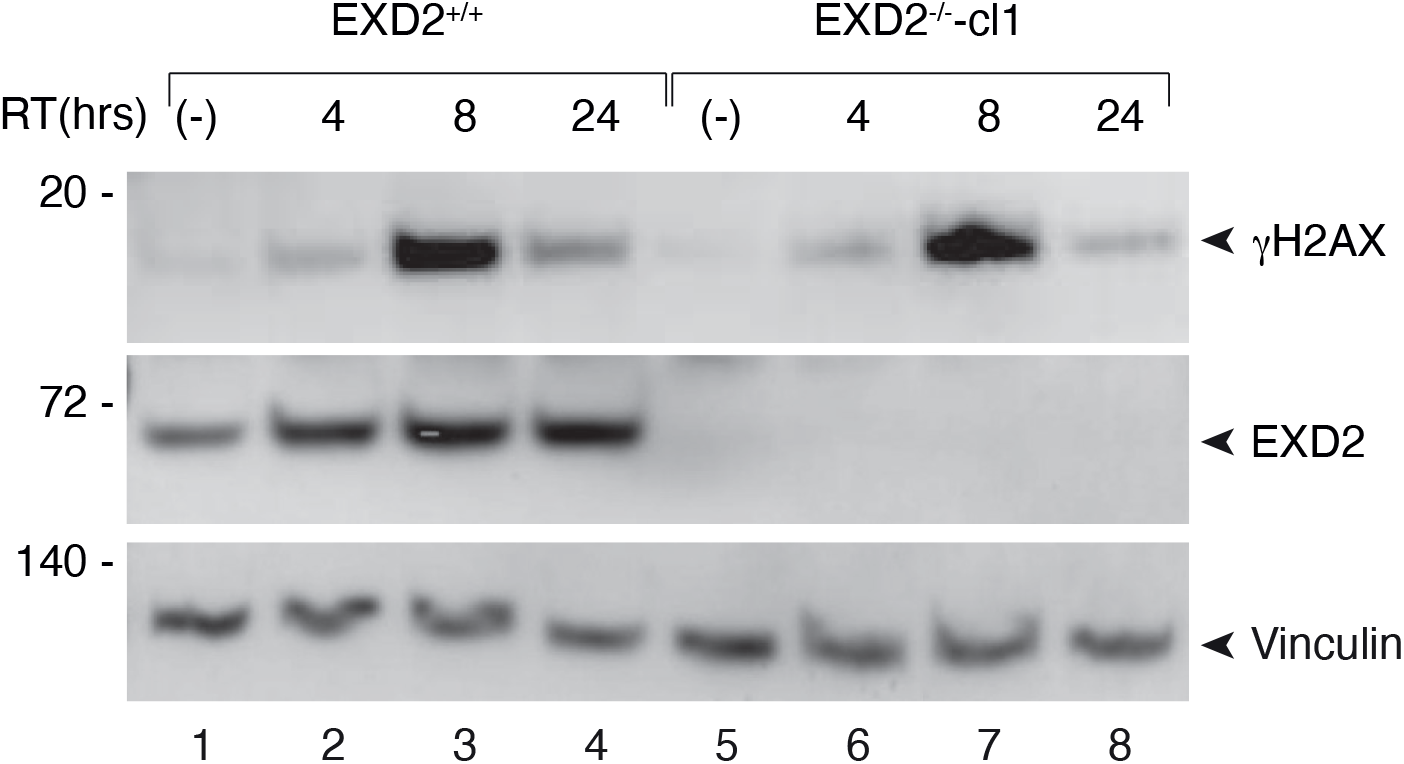
The DNA damage response is not affected by the lack of EXD2. EXD2^+/+^ and EXD2^-/-^-cl1 were mock or UV-irradiated (15J/m^2^) and let to recover for the indicated times before lysis. Protein lysates were immuno-blotted for EXD2, γH2AX or Vinculin. Molecular mass of the proteins is indicated (kDa). RT; recovery time.

**Supplemental Figure 5:**
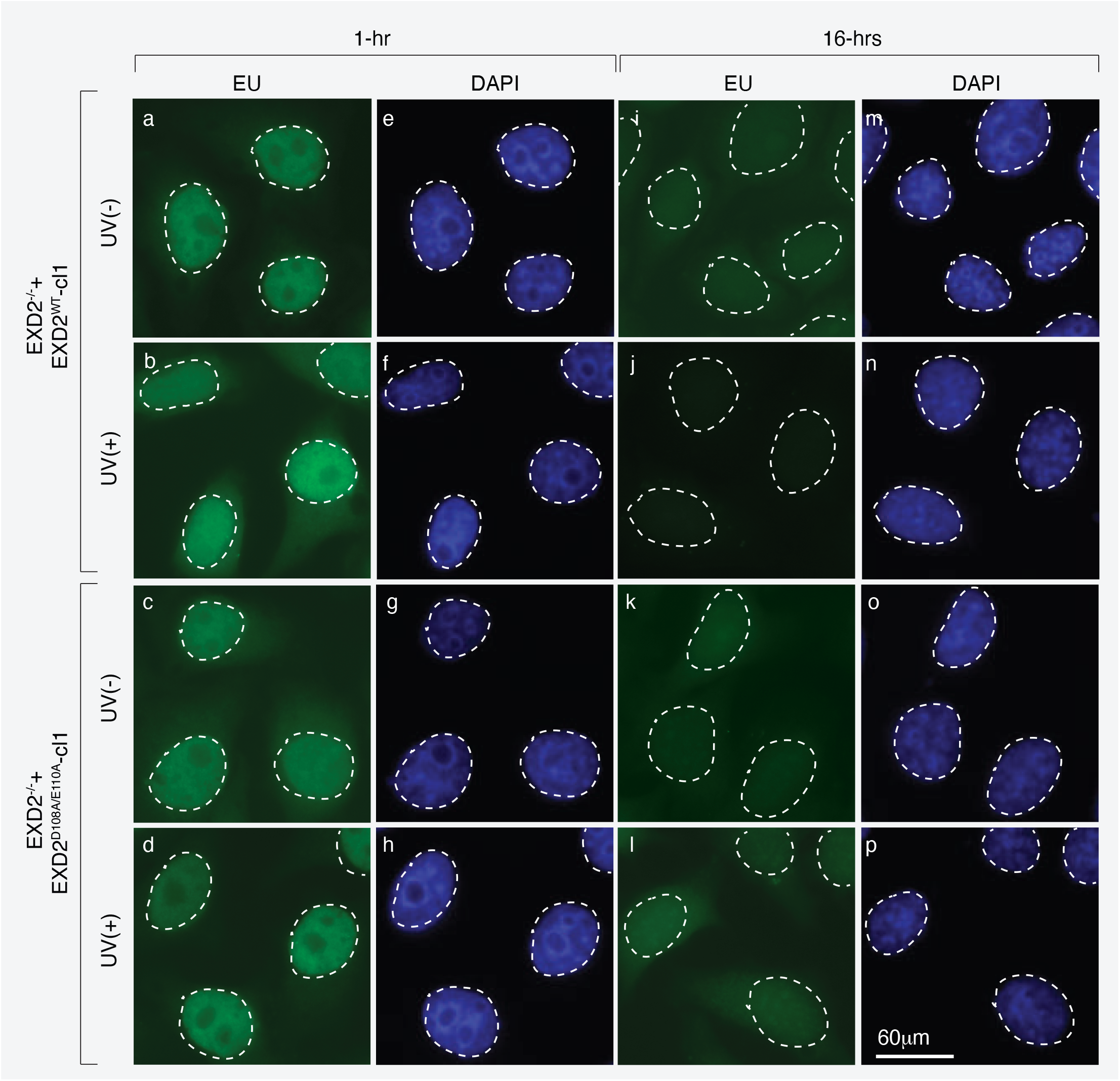
mRNAs under synthesis are degraded by EXD2 following UV irradiation. Representative confocal images of EXD2^-/-^ + EXD2^WT^-cl1 and EXD2^-/-^ + EXD2^D108A/E110A^-cl1 showing newly synthetized mRNA (1 hour or 16 hours after mock-treatment or UV irradiation at 15J/m^2^). Images of the cells were obtained with the same microscopy system and constant acquisition parameters.

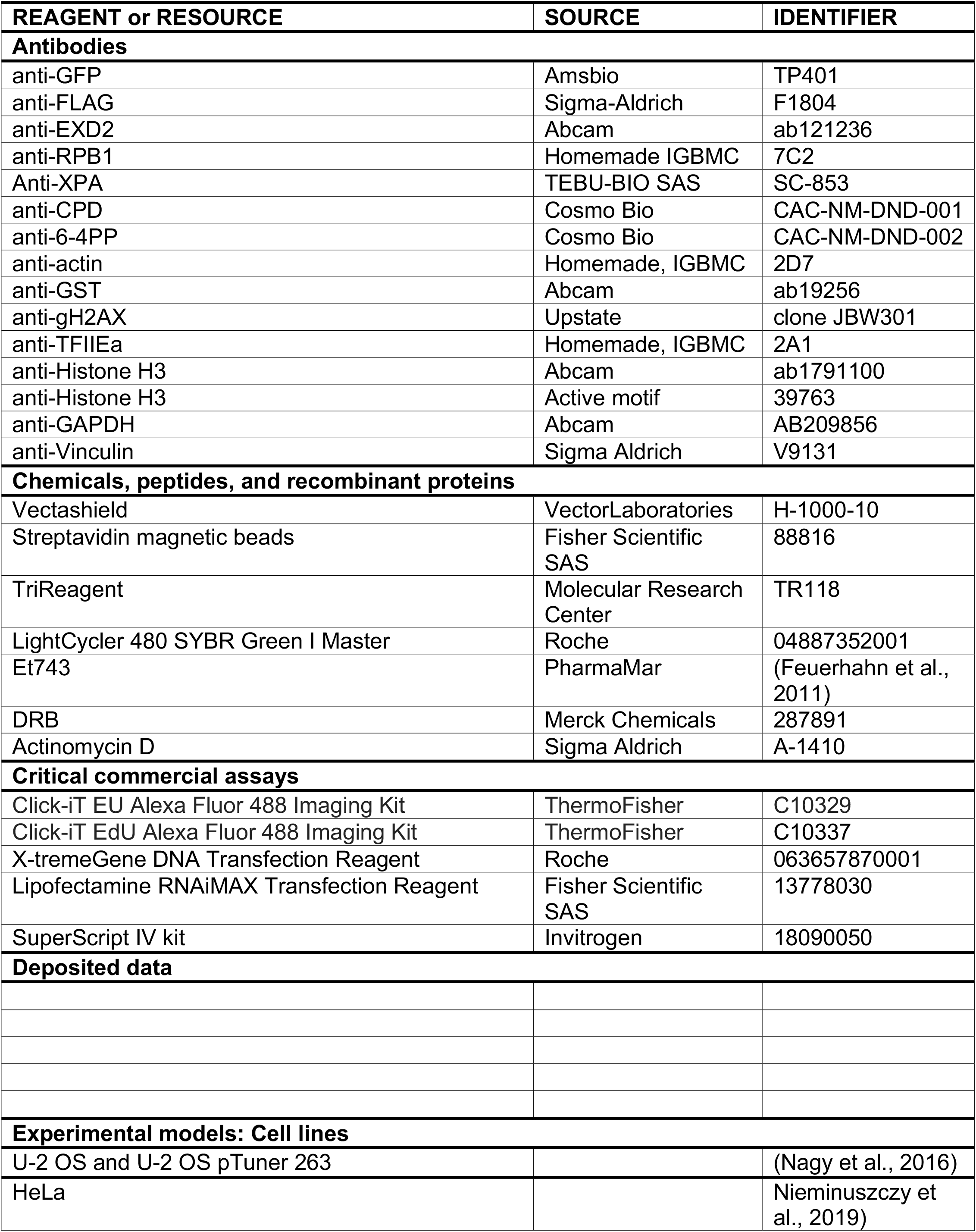
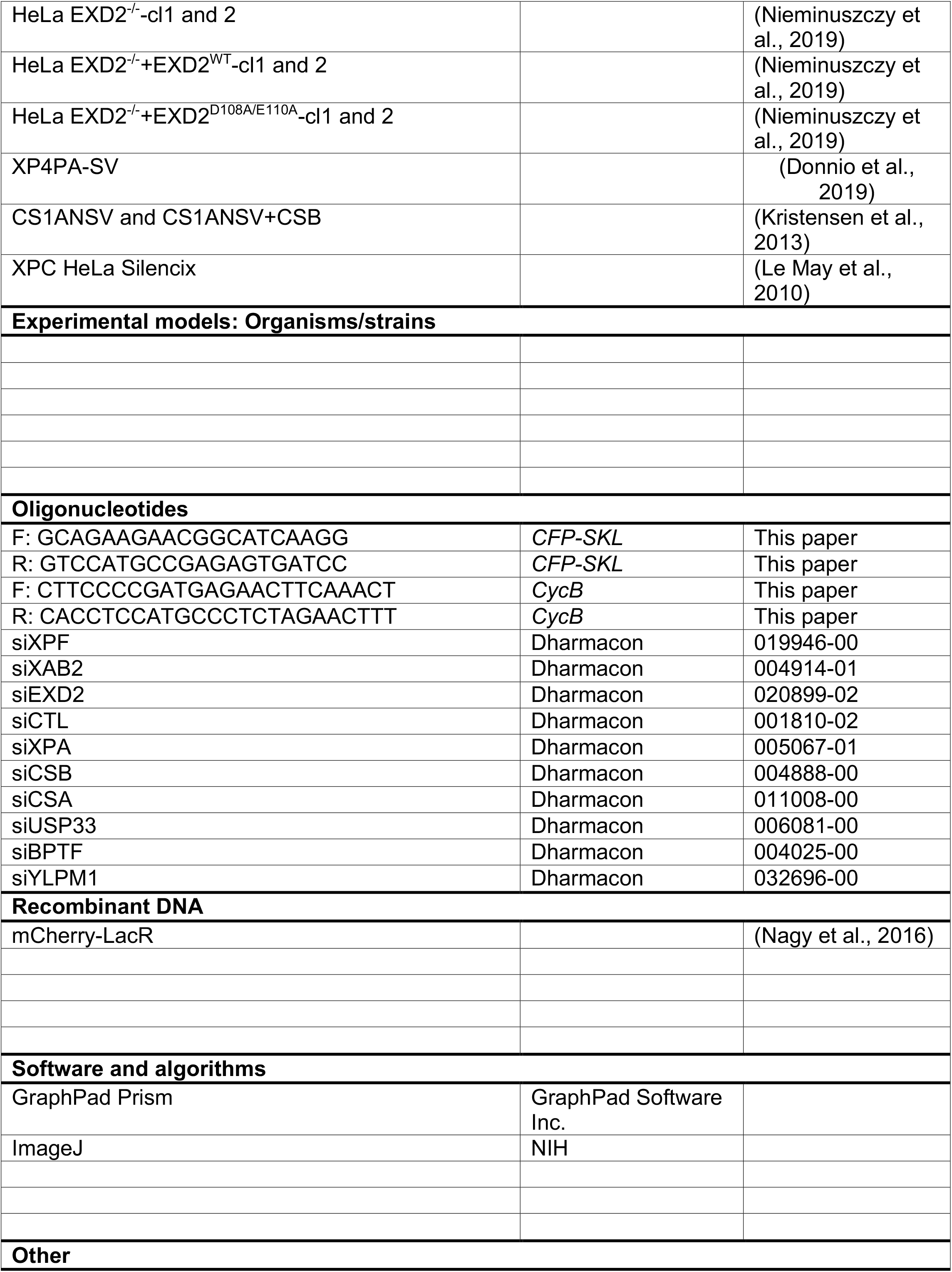
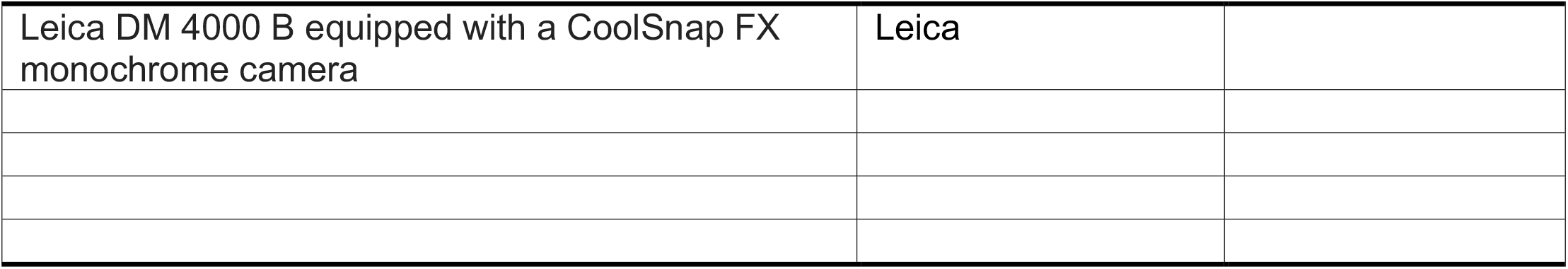

